# A convNet based multi label microRNA sub cellular location predictor, by incorporating k-mer positional encoding

**DOI:** 10.1101/2020.02.06.937656

**Authors:** Muhammad Nabeel Asim, Andreas Dengel, Sheraz Ahmed

**Author notes:** Correspondence: Muhammad Nabeel Asim, muhammad.

## Abstract

MicroRNAs are special RNA sequences containing 22 nucleotides and are capable of regulating almost 60% of highly complex mammalian transcriptome. Presently, there exists very limited approaches capable of visualizing miRNA locations inside cell to reveal the hidden pathways, and mechanisms behind miRNA functionality, transport, and biogenesis. State-of-the-art miRNA sub-cellular location prediction MIRLocatar approach makes use of sequence to sequence model along with pre-train k-mer embeddings. Existing pre-train k-mer embedding generation methodologies focus on the extraction of semantics of k-mers. In RNA sequences, rather than semantics, positional information of nucleotides is more important because distinct positions of four basic nucleotides actually define the functionality of RNA molecules. Considering the dynamicity and importance of nucleotides positions, instead of learning representation on the basis of k-mers semantics, we propose a novel kmerRP2vec feature representation approach that fuses positional information of k-mers to randomly initialized neural k-mer embeddings. Effectiveness of proposed feature representation approach is evaluated with two deep learning based convolutional neural network CNN and recurrent neural network RNN methodologies using 8 evaluation measures. Experimental results on a public benchmark miRNAsubloc dataset prove that proposed kmerRP2vec approach along with a simple CNN model outperforms state-of-the-art MirLocator approach with a significant margin of 18% and 19% in terms of precision and recall.

## 1 INTRODUCTION

Biological functionality of most of protein coding RNA is very much known, and explored rigorously [4]. There are substantial number of tools available which are capable to classify protein coding RNA [16, 6] and to predict protein sub-cellular localities [20, 2, 40]. Contrarily, biological functionalities of majority of non-coding RNAs (ncRNA) are still to be demystified [6]. Initially, non-coding RNAs were considered junk for a long period of time [33]. Nevertheless, with the progress of biological research, it was lately discovered that Non-coding RNA plays an important role in in multifarious noteworthy biological functions such as genomic imprinting, dosage compensation, and cell differentiation [6]. Non-coding RNA are also strongly associated with convoluted diseases such as Cancer, Alzheimer, and cardiovascular diseases [18, 32]. After such jaw-dropping findings, identifying new non-coding RNAs, and determining their biological functionalities quickly became hottest research area in Bioinformatics [6, 44].

Currently classification of non coding RNAs [6, 23] and prediction of non coding RNA sub cellular localization [43, 10] are topics of interest for many researchers. Non coding RNA sub-cellular localization plays an important role in gene regulation [13], neuronal dendrites [9], and embryonic development [25]. Almost in 80% human transcripts, asymmetrical localization usually happens because of delocalization of non coding RNA sequences at sub-cellular level [8]. Biological functions of non coding RNAs especially genetic information translation, transcriptional regulation and transduction of cellular signals are usually determined by the location of non coding RNA in specific cell, where every cell is primarily segregated into diverse compartments associated to distinct biological processes. For instance, miRNA localized to nuclear generally involves in mitosis or regulating gene expressions [11]. Likewise, sub-cellular localization of messenger RNA (mRNA) allows spatial and quantitative dominance over the production of proteins [43].

MiRNA sub-cellular localization is essentially required to regulate different scientific processes which usually take place within sub-cellular structures or organelles like mitochondrial metabolism carried by mito-miRNAs or synaptic plasticity carried by endosomal miRNAs [17]. MicroRNAs (miRNA) sub-cellular localization also largely contributes in several cellular processes of plants and animals involving development, differentiation, proliferation, and digestion in organisms along with post-transcriptional regulations of genes [3]. More recent research has discovered that few miRNAs also target epigenetic regulation function, and nucleus [26]. MiRNA sub-cellular localization facilitates proper interaction between RNA, and protein and it also decides miRNA action mode over target mRNAs. Although significant amount of time has been passed since the finding of miRNAs, however the way miRNAs regulate the gene repression is yet a point of contention [14]. Expanding evidences refers that mechanism behind gene silencing based on miRNA is extremely convoluted and can not be explained through a standalone model [21]. Deep analysis of how miRNAs manage to regulate their dedicated targets, one can acquire comprehensive understanding about the way cells accommodate gene expression processes to pathological and normal happenings of physiology at diverse scales of genetic information flow, and in several cellular compartments.

As the identification of sub-cellular localization of multifarious non coding RNAs (Lnc-RNA,mRNA, and miRNA) through biological experiments is highly labour intensive task and infallible to errors, hence development of computational methodologies for RNA sub-cellular location prediction is the need of an hour. Emergence of robust computational methodologies will accelerate RNA structural, and functional research enabling the practitioners to have a better picture of various biomedical implications. In order to explore the mechanisms behind RNA localization and deeply understanding biological functions of RNA, mainly the release of RNALocate metathesaurus [45] has played a significant role for the development of computational methodologies. RNALocate meta-thesaurus has over 37,700 entries for RNA sub-cellular localization along with experimental proofs such as 65 organisms (e.g Musmusculus, Homo sapiens, Saccharomyces cerevisiae), 42 sub-cellular localization’s (Endoplasmic reticulum, Nucleus, Cytoplasm, Ribosome) and 9 RNA classes (e.g miRNA, mRNA, lncRNA).

Through utilizing various sources such as RNALocate meta-thesaurus [45], ENCODE project [7], and Ensembl database [1], in last two years, three LncRNA [19, 36, 10], one mRNA [42] and one miRNA [42] sub-cellular localization classification methodologies have been proposed. While the performance of lncRNA [19], and mRNA [43] methodologies fall in 70s, miRNA methodology performance is even lower [42], in just 50s. In addition, predicting miRNA sub-cellular locality is way different than predicting locality for mRNA, and lncRNA sequences as miRNA sequences are quite shorter than mRNA, and lncRNA sequences. A very minor change in position, and length of nucleotides of a sequence may produce different location at sub-cellular level [42]. Moreover, according to the statistics of RNALocate Metathesaurus [45], almost 49% miRNA sequences may present in multiple compartments which indicate their ample localization patterns at sub-cellular level [42]. This indicates that miRNA sub-cellular localization is a multi-label classification problem. Because of these reasons, protein, mRNA, and lncRNA sub-cellular localization approaches can not be utilized for the prediction of miRNA sub-cellular localization.

Up to date, a limited amount of work has been performed for estimating the sub-cellular locality of miRNAs. One of the most eminent reason behind deficient work is the distinct sub-cellular localization properties of miRNAs, lack of prior knowledge-based features (e.g ontology), and their functional annotation in public datasets. According to our best knowledge, so far, there exists only one methodology for microRNA sub-cellular localization namely MIRLocator presented by Xiao et al. [42]. MIRLocator utilized a sequence-to-sequence model which used non-arbitrary label order. In Natural Language processing (NLP), several researchers such as Vinyals et al [39] have proved that label order has a huge effect on the generalization ability of sequence-to-sequence model. However, as label order is pre-defined in MIRLocator, so the performance is significantly depending on prior information related to label order. In addition, even the model manages to predict all true labels accurately, irrational training loss may still occur because of inconsistent order of the labels. Thus, it can be summarized that model performance is highly sensitive to pre-declared order of labels. However, other deep learning models which do not use sequence to sequence approach, for them any label order shall work effectively without considering pre-declared label order information.

Most of the existing DNA and RNA sequence analysis approaches generally rely on k-mers of DNA or RNA sequences which are generated by sliding a fixed size window over the sequences with a particular stride size [42, 37, 34]. As K-mers are just chunk of characters, thus, they are usually treated as standard words in Natural Language Processing (NLP) [6]. Considering the similarity of k-mers with textual data, and inspiring from the performance of pre-trained neural word embeddings in NLP, many researchers have developed pre-trained neural k-mer embeddings for various bio-infomatics tasks like prediction of Chromatin accessibility using 6-mers with glove embeddings [28], protein retrieval and sequence classification using seq2vec and prot2vec [22], and Glove based k-mer embeddings for micro RNA sub cellular localization [42]. However, *K – mer* based pre-trained embeddings do not produce significant improvement in the performance of DNA or RNA sequence analysis as standard neural word embeddings have produced in diverse NLP tasks [12].

Publicly available approaches for the learning of word embeddings from textual data, DNA or RNA sequences operate on a basic principle in which a fixed size window is convolved on sequences or textual data [15, 29]. In a fixed size window, semantic information of words is extracted on the basis of their surrounding words. It is relatively easy to capture semantic information of words in natural language processing as compare to capturing the semantics of k-mers in DNA or RNA sequences. Mainly four basic nucleotides (A,G,C,T or U) encode the grammatical information of DNA or RNA sequences where distinct positions of these nucleotides actually define the functionality of the sequences. Considering the dynamicity of nucleotides positions, instead of learning representation on the basis of k-mers semantics, we utilize positional information of k-mers to generate k-mer embeddings of DNA or RNA sequences.

Furthermore, performance of deep learning model varies at different k-mers due to difference in distribution of basic four nucleotides (A,C,G,T or U) [24]. As it is a tedious task to generate pre-trained embeddings of different k-mers, hence rather than utilising pre-trained neural k-mer embeddings, we propose a novel light weight kmerRP2vec feature representation approach. The proposed kmerRP2vec approach precisely captures the positional information of k-mers in miRNA sequences. It first encodes positional information of each k-mer into a fixed length vector. Then, encoded positional information is fused into randomly initialized k-mer embeddings. To predict micro RNA sub-cellular locations, we propose a convolutional neural network based approach named as “MirLocPredictor” which neither require any information of pre-declared label order nor pre-trained k-mer embeddings. Also, in order to evaluate the performance impact of proposed feature representation approach on a recurrent neural network based approach, we adapt a classification methodology namely TextRNN proposed by Liu et al [27]. TextRNN has produced state-of-the-art performance for text document classification.

Finally by using both proposed CNN based methodology and adapted RNN methodology, a fair performance comparison of proposed kmerRP2vec, and 4 other feature representation approaches is performed using 8 renowned evaluation measures. Experimental results have proved that proposed feature representation approach has significantly improved the performance of both classification methodologies. Overall, proposed MirLocPredictor substantially outperforms both adapted TextRNN and state-of-the-art MIRLocator methodologies.

Our contribution can be summarised as follows:

1. We propose a novel kmerRP2vec feature representation approach which captures the positional information of the k-mers of miRNA sequences. This positional information is injected with randomly initialized neural k-mer embeddings.
2. We perform extensive experimentation with two diverse classification methodologies to prove the effectiveness of proposed feature representation approach.
3. We present a simple end to end system for the prediction of miRNA subcelular localization.

## 2 MATERIALS AND METHODS

This section briefly describes proposed MirLocPredictor and adapted TextRNN methodologies. It also discusses the characteristics of experimental dataset followed by performance evaluation measures.

### 2.1 Proposed Methodology

In Natural Language Processing, following the success of transformer [38] based approaches, here for the very first time we capture the positional information of k-mers from miRNA sequences and fuse this positional information in randomly initialize k-mer embeddings. Furthermore, we use one dimensional convolutional layer for the extraction of discriminative features from k-mers of miRNA sequences. As miRNA sequences are very small in size, so descriminative features extracted by convolutional layer along with neural embedding features are concatenated before passing to two fully connected layers. The final fully connected layer is used as a classifier for the prediction of various locations associated with the miRNA sequences. Figure 1 shows graphical representation of proposed MirLocPredictor methodology. Subsequent sections briefly describes main modules of miRNA sub cellular localization methodology.

**Figure 1.**
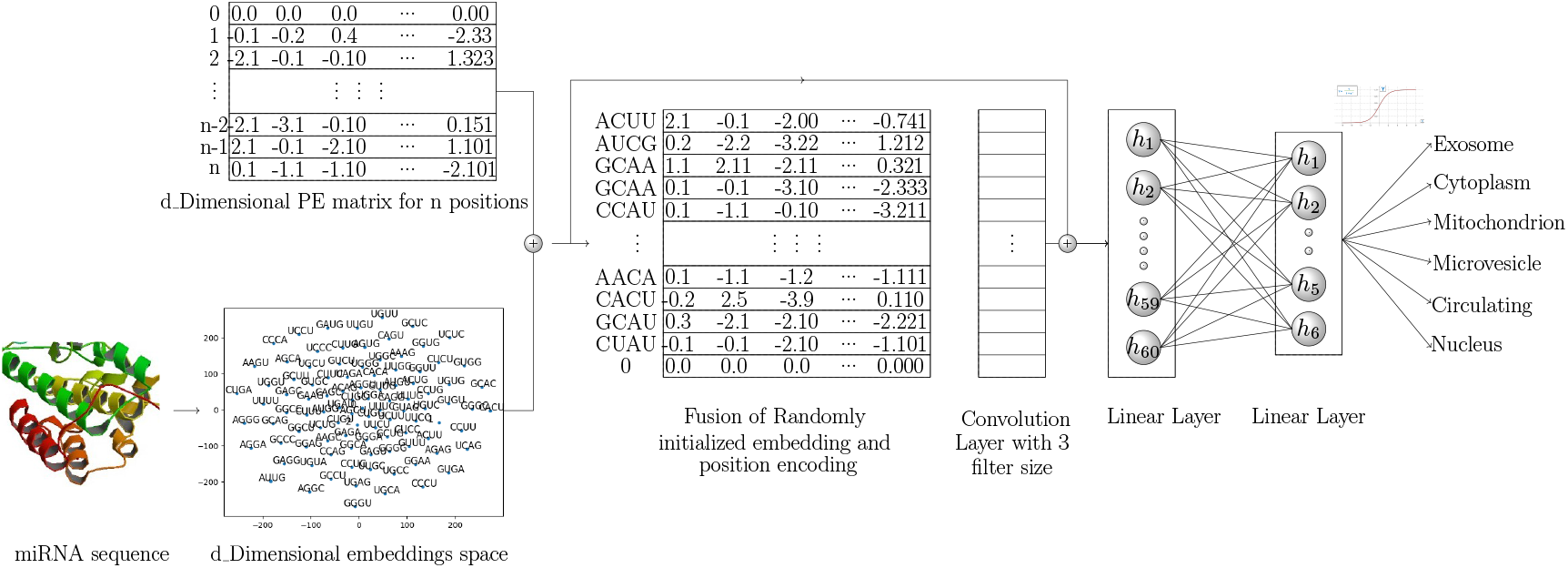
Graphical Representation of proposed MirLOcPredictor.

#### 2.1.1 Feature Representation

Proposed kmerRP2vec feature representation approach consists of three steps. Firstly, we randomly generate k-mer embeddings. Secondly, we capture the positional information of all k-mers in a sequence. The positional information is encoded to a fixed length vector equal to the length of randomly initialize embeddings. Finally we aggregate vectors of randomly initialize embeddings and positional encoded vectors to get final representation of each k-mer. A pseudo code to generate Positional encoding is given below:

**Algorithm 1:**
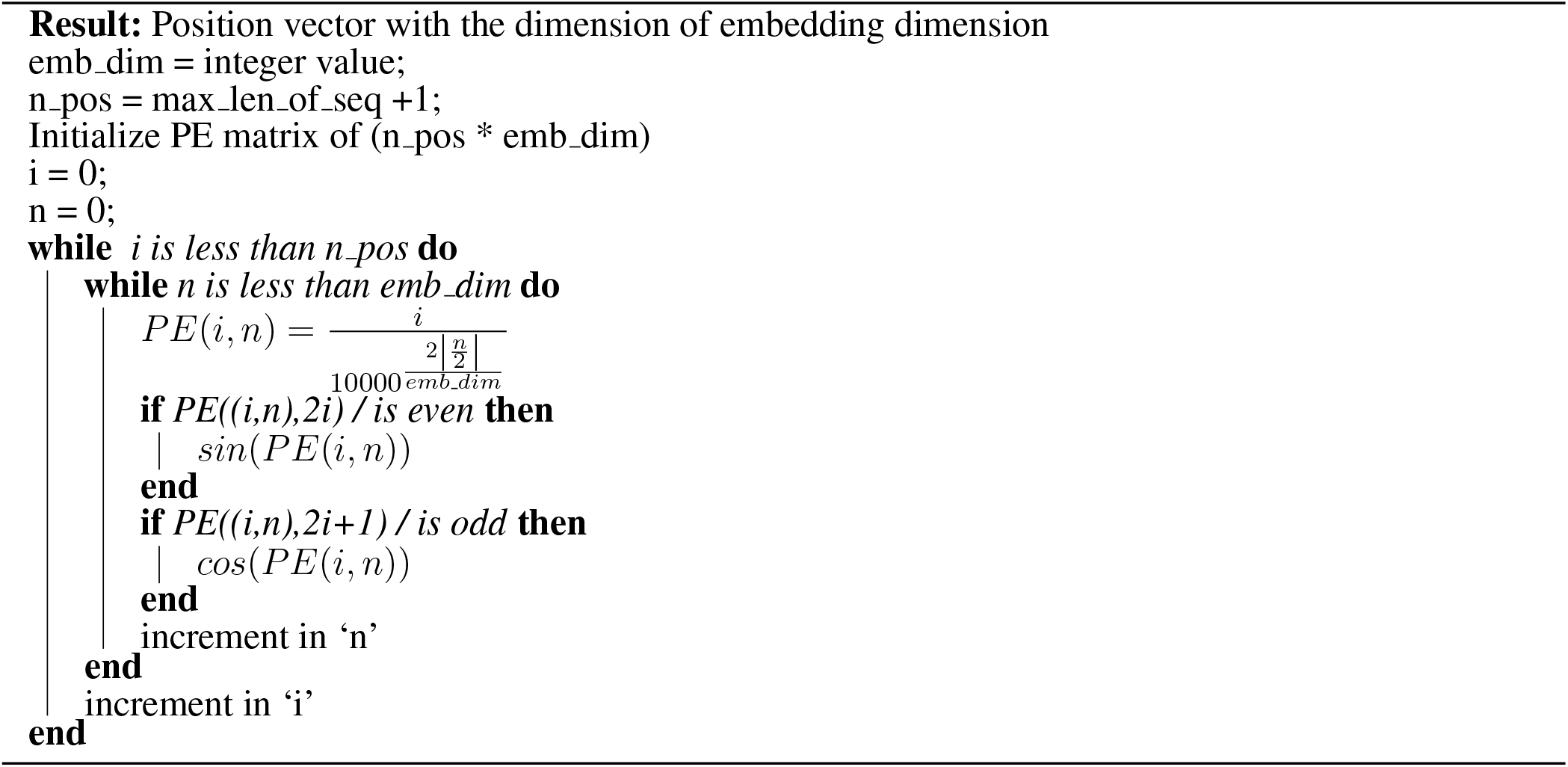
Pseudo code to create Positional Encoding

In Pseudo code, outer loop shows total number of positions for unique k-mers in the miRNA subcellular localization dataset, and inner loop represents the dimension of positional encoding. Rather than denoting the position of k-mers with binary values which consume more memory, we have encoded positional information using sine and cosine functions. These functions represents the alteration in positional bits in such a way that even and odd position values are in the range of sin(x) and cosine(x) respectively. This representation is also known as sinusoidal representation where the range of all Real numbers ℝ is fixed from −1 to 1. Thus it provides unique encoding even for long sequences at every time step. Sinusoidal position encoding makes symmetrical distance between neighboring time-steps and allows representation decay nicely with time.

For Positional encoding, matrix maximum length of sequence is fixed by taking the maximum length of miRNA sequence. Sequences which have length greater than max len seq are trimmed, and sequences with smaller length are padded with zeros. PE[0] vector represents the all locations in sequences which are padded by zeroes, while PE[1] represents k-mer at first index in each sequence similarly PE[2] for k-mer at second index and so on. To get final representation, we have fused positional encoded vectors with randomly initialized embedding vectors of all k-mers.

#### 2.1.2 Feature Extraction

As miRNA sequence data is one dimensional so we utilise a one dimensional convolution layer for the extraction of discriminative features. Suppose miRNA sequence has n length k-mers *Seq*_1:*n*_ = *k*_1_, *k*_2_,…, *k_n_*,, where each k-mer is associated with *d_dimensional* embedding vector. Over all sequences, convolution is performed with filter size of *k* = 3, stride size 1 which produces 1D convolution of *width-k*. In order to add non-linearity into this, 1D convolution is passed through Relu activation function. Mathematically convolutional process can be expressed as follows:

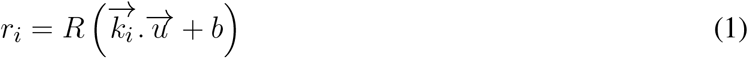

Where 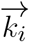 is d dimensioal embedding vector of *i_th_* index kmer, 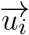 is weight matrix vector, b is bias value and R represents Relu activation function.

### 2.2 TextRNN

While CNNs are good at attaining position invariant and local features, RNNs are considered more appropriate to capture long range contextual information. To evaluate the performance impact of proposed feature representation approach on the performance of recurrent neural networks. Here we adopt Liu et al. [27] text classification methodology. Adopted methodology has lstm layer followed by attention layer which helps to extract more discriminative and focused features.

### 2.3 Data set

In order to evaluate the integrity of proposed methodology for the task of miRNA sub-cellular localization, we use a publicly available benchmark data set provided by Xiao at al [42]. Originally, entire data of microRNA sub-cellular localization was collected from RNALocate metathesaurus [45], and miRNA sequences were gathered from miRBase repository 1. Provided dataset has 1047 miRNA sequences which are annotated against the combination of 6 cellular locations namely Exosome, Cytoplasm, Mitochondrion, Microvesicle, Circulating, and Nucleus. Furthermore, each sequence is made up of 4 nucleotides A, C, U, G and overall sequence length lies between 20/30 nucleotides. Further statistics of benchmark dataset against each class label are summarized in Table 1.

**Table 1.** Characteristics of Benchmark MiRNA Sub-Cellular Localization Dataset [42]

| MiRNA Location Distribution |  |  |  |  |  |
| --- | --- | --- | --- | --- | --- |
| Exosome | Cytoplasm | Mitochondrion | Microvesicle | Circulating | Nucleus |
| 869 | 209 | 338 | 348 | 513 | 349 |
| MiRNA Class Distribution |  |  |  |  |  |
| Uni-Label | Bi-Label | Tri-Label | Tetra-Label | Penta-Label | Hexa-Label |
| 424 | 233 | 128 | 78 | 64 | 120 |

### 2.4 Performance Measures

Evaluation of multi-label classification methodologies is quite difficult and a way different task than evaluating multi-class classification methodologies [30, 41]. In multi-class classification prediction can be either fully correct or incorrect, however in multi-label classification, prediction can be fully correct, incorrect or partially correct [20]. Evaluation of multi label classification methodologies is considered similar to the evaluation of information retrieval methodologies. In order to evaluate the performance of miRNA sub-cellular location prediction methodologies, we use 8 different evaluation measures which have been widely used to evaluate the performance of information retrieval and multi-label text classification methodologies.

Let suppose, *C* is a multi-label corpus consists of |*C*| number of multi-label examples where each example is represented as (*a_i_, B_i_*), i = 1,2,3,… |*C*|, *B_i_* ⊆ *L*. Let H is a classifier that predicts the label set *Cl_i_* for instance *a_i_*, where predicted label set is represented as Pl.

#### 2.4.0.1 Accuracy

In binary or multi-class classification accuracy is computed by taking ratio of predicted correct labels to the total number of labels. However, to evaluate the performance of multi-label classification methodologies, for each sequence sample, we compute the ratio between correctly predicted labels and total number of labels (predicted and actual labels)[5]. Overall accuracy is computed by taking the average across all instances of the dataset. Mathematically, it can be expressed as:

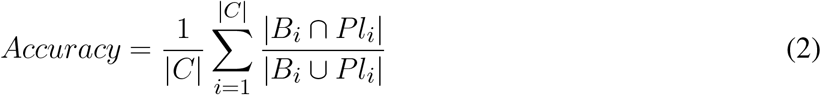

#### 2.4.0.2 Precision

Precision of a sample sequence is computed by taking ratio between correctly predicted labels and actual labels of the particular sequence sample [5]. Finally, overall precision is computed by taking the average across all instances of the dataset. Mathematically precision can be defined as:

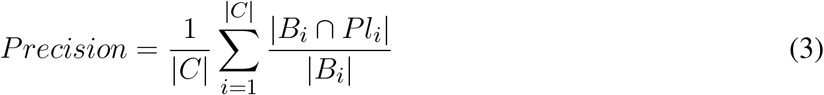

#### 2.4.0.3 Recall

Recall is a proportion of correctly predicted labels to overall predicted labels of a sample sequence [5]. Recall of all samples is calculated by averaging the recall across all samples of the corpus. Its mathematical formula can be expressed as:

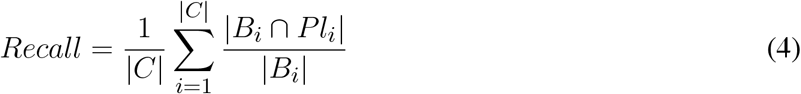

#### 2.4.0.4 Average Precision

For every relevant label, it estimates how many relevant label are actually ranked before it, and takes mean against set of relevant labels. Mathematically, average precision can be defined as:

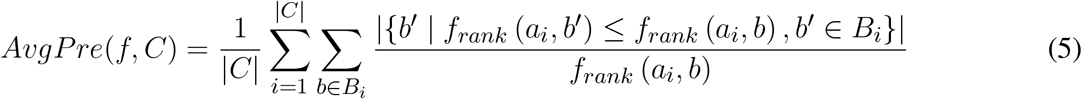

To evaluate the indignity of proposed methodology, the idea of average precision is borrowed from Information Retrieval, where it has been extensively used to evaluate the ranking of relevant documents retrieved against certain query [31]. Average precision is directly proportional to the performance of the model.

#### 2.4.0.5 F_1_-Measure (F)

*F*_1_-Measure is the harmonic mean between *Precision* and *Recall* [5]. In multi-label classification definitions of recall and precision leads to the following definition of *F*_1_-Measure:

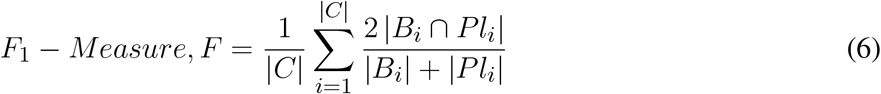

#### 2.4.0.6 Micro F_1_-Measure (F)

Harmonic mean among micro-precision, and micro-recall refers to Micro-F1 which cane be defined as:

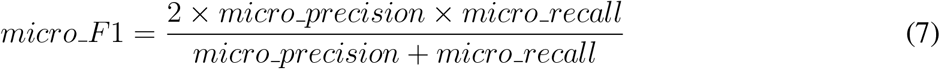

#### 2.4.0.7 Macro F1-Measure (F)

Macro-F1 is the harmonic mean among trivial multi-label precision, and recall where first of all average is computed for single sample sequence and afterwards mean across all corpus is taken [42].

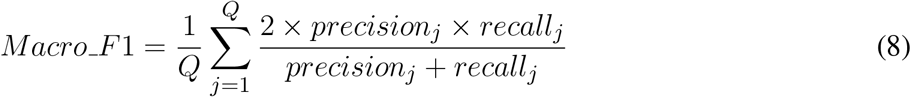

#### 2.4.0.8 Hamming Loss (HL)

Hamming loss estimates the frequency to which a sample-label is incorrectly classified and it mainly focus on the labels which are not predicted at all (missing the prediction of a relevant label) or are wrongly predicted (prediction error). Mathematically hamming loss [35] is defined as:

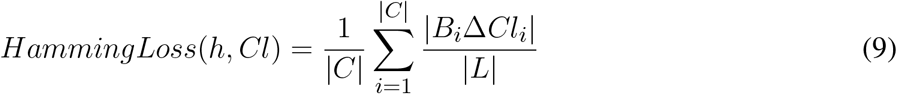

In this equation, delta (Δ) shows the symmetric difference between two sets(*B*i and *Cl_i_*). In other words, delta (Δ) acts like XOR operation in the Boolean logic. Hamming loss predicts to what extent actual and predicted labels are dissimilar. Hamming loss of zero means that a classifier has predicted all the labels accurately, whereas higher than zero value depicts that prediction is not error-free. So from this, it is easy to deduce that hamming loss and accuracy are inversely proportional.

## 3 EXPERIMENTAL SETUP AND RESULTS

In order to generate different k-mers of miRNA sequences, we slide a window of different sizes on the sequence with the same stride size of 1. Furthermore, we implement presented positional encoding based feature representation approach in python and proposed deep learning model is implemented in Pytorch.

Following Xiao et al. [42], we have preformed 10-fold cross validation where at each fold one part of data is utilised for testing, from other 9 parts, 10% data is used for validation, and remaining 90% data is utilised to train the model.

### 3.1 Results

To prove the effectiveness of proposed kmerRP2vec feature representation approach, we have compared its performance with 4 different feature representation approaches using 2 distinct deep learning based classification models. Firstly, we feed the deep learning methodologies with randomly initialized 120 dimensional word vectors followed by pre-trained neural k-mer embeddings provided by Xiao et al. [42]. Thirdly, we generate neural k-mer embeddings solely based on the positional information of each k-mer. To reap the benefits of both pre-trained and positional neural k-mer embeddings, we also perform experimentation by combining both of these embeddings. Finally, we utilize a novel kmerRP2vec feature representation approach which injects positional information of every k-mer with randomly initialized neural k-mer embeddings.

All feature representation approaches are named with appropriate prefixes such as pre-trained neural k-mer embeddings are referred as pre-embedding, randomly initialise embeddings are named as Rand-Embedding, position based embeddings are named as Pos-Encodding, and positional information fused with pre-trained neural k-mer embeddings are refferd as Pre-Embedding + Pos-Encodding.

#### 3.1.1 Performance Impact of Proposed Feature Representation Approach

This section illustrates the performance impact of 5 different feature representation approaches on the performance of proposed CNN based methodology “MirLocPredictor” and adapted TextRNN [27] based methodology.

Table 2 reports the performance figures of MirLocPredictor, and adapted TextRNN [27] methodology produced using 5 different feature representation approaches in terms of 8 evaluation metrics.

**Table 2.**
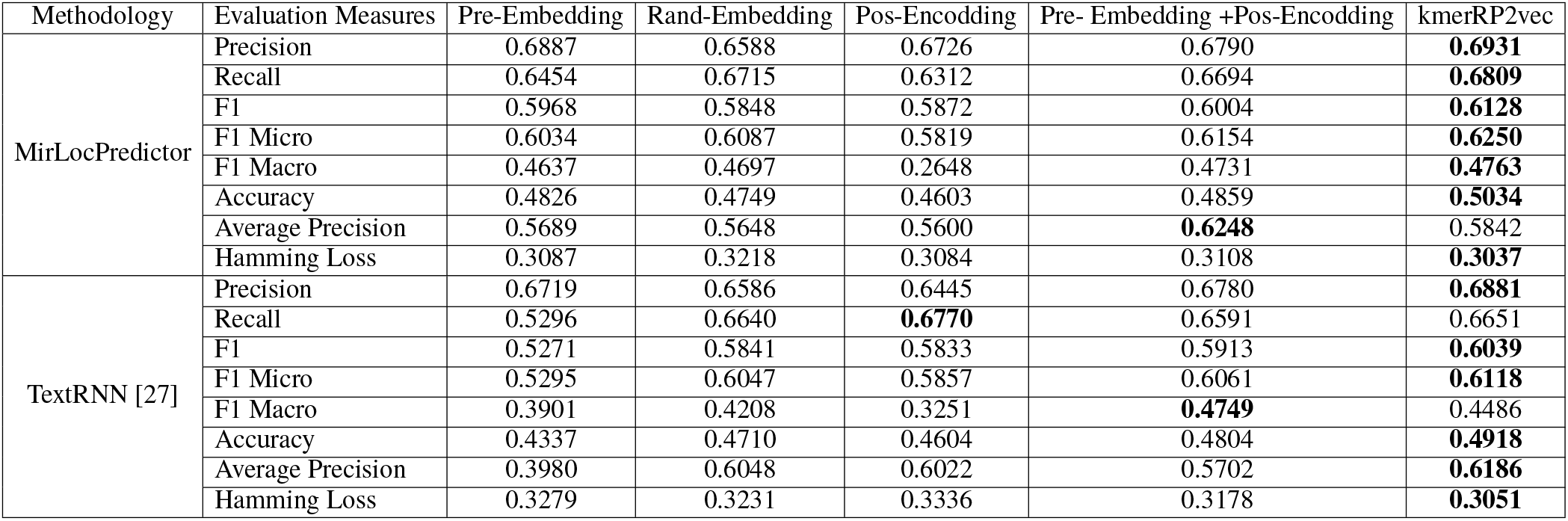
Performance comparison between proposed kmerRP2vec and 4 other feature representation approaches on account of 4-mers using two classification methodologies

As is depicted by the Table 2, MIRLocPredictor marks almost similar performance with randomly initialized and pre-trained neural k-mer embeddings. Because both embeddings produce better performance on 4 distinct evaluation metrics. Moreover, amongst all feature representation approaches excluding proposed one, MIRLocPredictor manages to achieve highest precision of 69% using pre-trained neural k-mer embeddings, highest recall of 67% is achieved using randomly initialized k-mer embeddings. Although MIRLocPredictor marks lowest hamming loss with the use of position encoding embeddings, however, overall MIRLocPredictor performance declines even lower than randomly initialized and pre-trained neural k-mer embeddings when evaluated across other performance measures. From all k-mer embeddings excluding proposed feature representation approach, hybrid approach (Pre-embedding + Pos-Encoding embeddings) produces better performance across 5 evaluation metrics (F1, F1-micro, F1-macro, Accuracy, Average Percision). On the other hand, amongst all feature representation approaches MIRLocPredictor produces most promising performance using proposed kmerRP2vec feature representation approach across 7 evaluation metrics. Only for Average Precision measure MIRLocPredictor produces better performance when it is fed with hybrid feature representation (Pre-Embedding + Pos-Encodding) approach. In a nutshell, it can be concluded that MIRLocPredictor has better performance with both feature representation approaches Pre-Embedding + Pos-Encodding and kmerRP2vec as compared to the performance produced using simple pre-trained or randomly initialized k-mer embeddings.

Turning towards other half of the Table 2, adapted TextRNN produces superior performance across 7 evaluation metrics using randomly initialized neural k-mer embeddings as compared to pre-trained and position encoding based embeddings. Amongst all feature representation approaches excluding proposed approach, as similar to MIRLocPredictor, adapted TextRNN approach [27] produces top performance figures with hybrid embedding approach (Pre-embedding and Pos-Encoding embeddings). Hybrid (PreEmbedding + Pos-Encodding) approach outshines other three feature representation (Pre-Embedding, Rand-Embedding, Pos-Encodding) approaches over 6 evaluation metrics (Precision, F1, F1-micro, F1-macro, Accuracy, Hamming Loss). Contrarily, once again, proposed kmerRP2vec approach outperforms all other feature representation approaches by significantly increasing the performance of adapted TextRNN methodology. To sum up, it can be concluded that proposed feature representation approach (kmerRP2vec) significantly improves the performance of both classification models.

It is a tedious task to generate pre-trained neural embeddings for different k-mers. As shown by the Table 2, amongst all feature representation approaches excluding proposed one, only randomly initialized neural k-mer embeddings manages to produce comparable performance to pre-trained neural k-mer embeddings with both deep learning based classifiers. Thus, for further experimentation we have compared the performance of randomly initialise k-mer embeddings with proposed kmerRP2vec feature representation approach. Figures 2 and 3 illustrate the performance of MirLocPredictor and Adapted TextRNN [27] over both randomly initialized neural k-mer embedding, and proposed kmerRP2vec feature representation approach using 8 different k-mers. In order to make the graphs thoroughly visualizeable, we have mapped the results of only 4 evaluation metrics (F1, F1-macro, F1-micro, Average Precision) in the aforementioned Figures. For better understanding and to make graphs more readable, the performance of both classification models using randomly initialized neural k-mer embedding is named with the prefix of evaluation metrics (e.g f1,P, R) followed by R. Similarly performance with proposed kmerRP2vec feature representation approach is named with the prefix of evaluation metric followed by kmerRP2vec. Here R shows Randomly initialise k-mer embeddings and kmerRP2vec represents Positional encoded + Randomly initialised k-mer embeddings.

**Figure 2.**
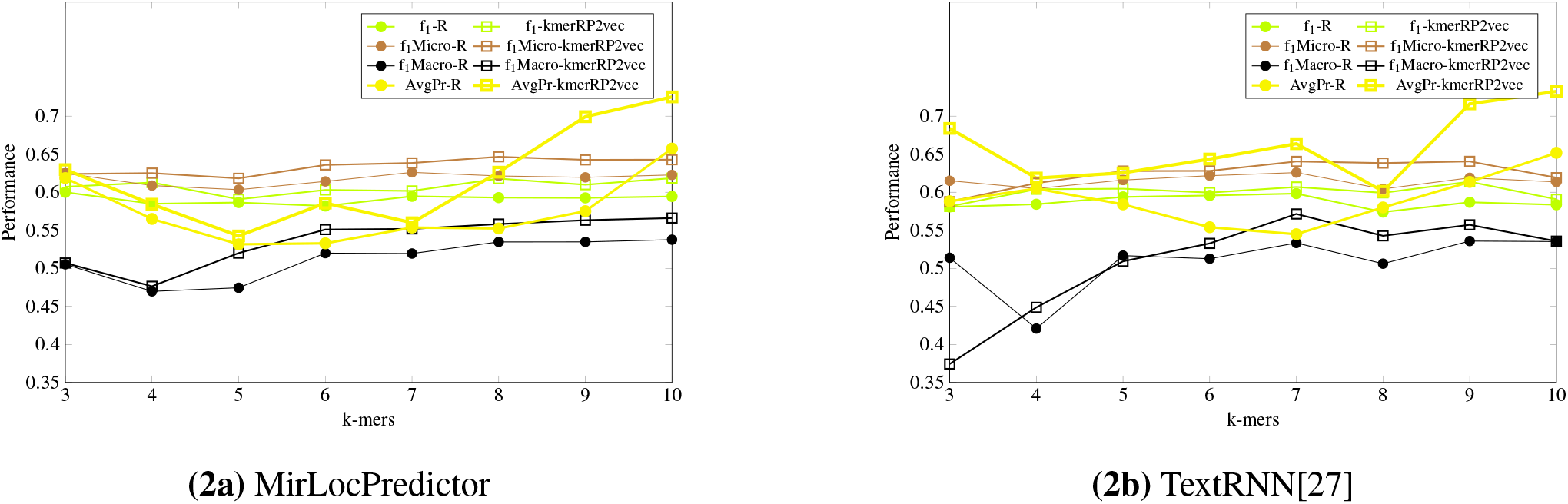
Performance comparison of proposed kmerRP2vec with randomly initialized k-mer embeddings at 8 benchmark k-mers using two classification methodologies

**Figure 3.**
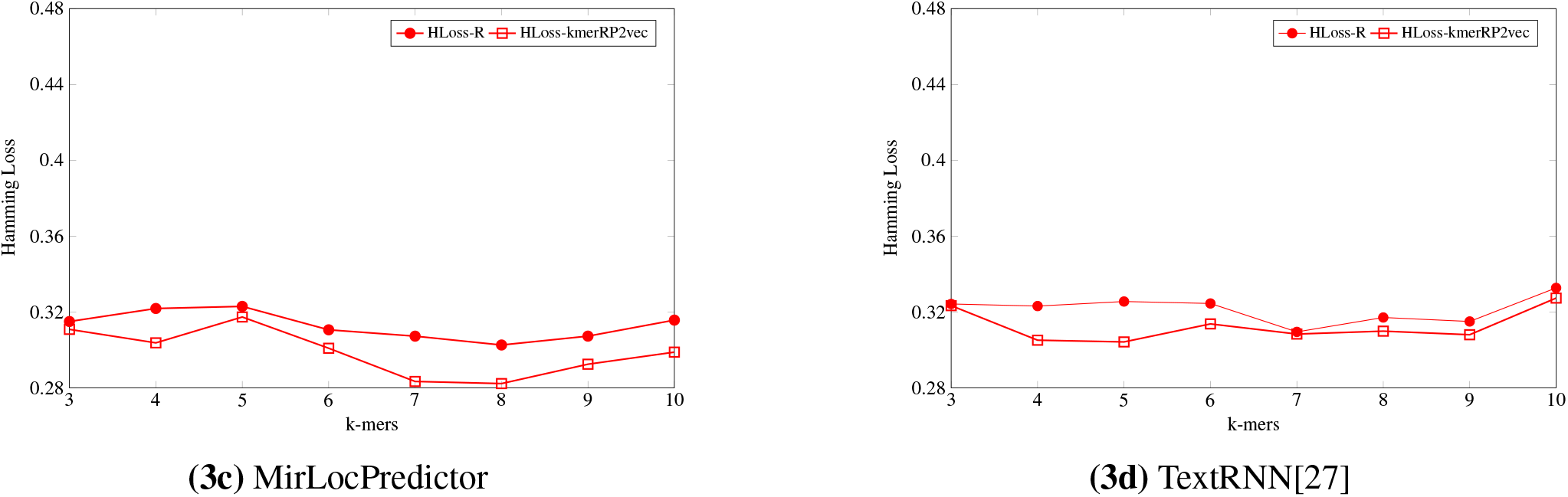
Hamming Loss of proposed kmerRP2vec and randomly initialized k-mer embeddings at 8 benchmark k-mers using two classification methodologies

As is shown by Figure 2, in terms of *F*_1_ evaluation measure, both classification models perform better with proposed kmerRP2vec feature representation approach by marking the higher performance almost across all k-mers. Randomly initialised k-mer embedding only manages to equalize the promising performance of proposed kmerRP2vec twice (k-mers:5,7) with MirLocPredictor and thrice (k-mers:5,6,7) with adapted TextRNN. Likewise, taking F1 variants into account (micro, macro), although performance of MirLocPredictor computed using proposed kmerRP2vec feature representation approach initially remains close to the performance produced by randomly initialized neural k-mer embeddings until 4-mers as compared to 5-mers for TextRNN. However, afterwards, proposed feature representation approach significantly increases the performance of both classification methodologies. Just like F1-score, average precision produced using proposed kmerRP2vec feature representation approach remains very high at majority of k-mers for both MirLocPredictor, and TextRNN methodologies.

On the other hand as sown by Figure3, evaluating the performance of both classification methodologies in terms of hamming loss, MirLocPredictor hamming loss using proposed kmerRP2vec feature representation approach is significantly lower across all k-mers than the hamming loss produced using random initialised k-mer vectors. Whereas, TextRNN produces same hamming loss values with both feature representation approaches only at two k-mers (3, 7), however, at most k-mers TextRNN hamming loss using proposed kmerRP2vec feature representation approach also remains lower than the hamming loss produced by randomly initialized k-mer embeddings.

To summarize, across all evaluation metrics, proposed kmerRP2vec feature representation approach significantly raises the performance of both classification methodologies.

#### 3.1.2 Performance Comparison of Proposed, Adapted, And State-of-the-Art MiRNA sub-cellular location prediction Methodologies

This section compares the performance of proposed MirLocPredictor with adapted TextRNN and state-of-the-art MirLocator methodologies for the task of miRNA sub-cellular location prediction.

Table 3 reports the performance figures produced by two classification methodologies namely MirLocPredictor and TextRNN [27] using 8-mers, and performance figures of MirLocator [42] using 4-mers in terms of 8 different evaluation metrics. As the Table 3 suggests, adapted TextRNN approach shows better performance as compared to state-of-the-art MirLocator across all evaluation metrics. However, TextRNN approach seems more biased towards type 1 error as it has high precision and low recall. On the other hand, amongst all, proposed MirLocPredictor significantly outperforms state-of-the-art MirLocator approach across all evaluation measures. Comparing the performance values of proposed and adapted methodologies, MirLocPredictor is a clear winner as it performs better across 7 evaluation measures compared to TextRNN which only manages to mark higher recall values. Hence, overall performance of proposed MirLocPredictor approach is better amongst all as it is neither biased towards type 1 nor type 2 errors.

**Table 3.** Performance comparison of proposed MirLocPredictor with both state-of-the-art MIRLocator [42] and adapted TextRNN[27] Approaches

| Methodology | Evaluation Measures |  |  |  |  |  |  |  |
| --- | --- | --- | --- | --- | --- | --- | --- | --- |
|  | Precision | Recall | F1 | F1 Micro | F1-Macro | Accuracy | Average Precision | Hamming Loss |
| TextRNN[27] | 0.6237 | **0.7092** | 0.5992 | 0.6383 | 0.5427 | 0.4773 | 0.6012 | 0.3099 |
| MIRLocator[42] | 0.5033 | 0.4849 | – | 0.5820 | 0.4933 | – | 0.5820 | – |
| MirLocPredictor | **0.6878** | 0.6784 | **0.6178** | **0.6465** | **0.5581** | **0.5051** | **0.6263** | **0.2822** |

### 3.2 Conclusion

This paper proposes a novel kmerRP2vec feature representation approach which fuses positional information of k-mers and randomly initialized k-mer embeddings. Through precisely analysing the performance of different feature representation approaches with two distinct classification methodologies across 8 evaluation measures, we concluded that pre-trained neural k-mer embeddings are not producing promising performance in miRNA sequence analysis tasks similar to what neural word embeddings managed to produce in Natural Language Processing tasks. Primarily, this is because of the fact that in DNA or RNA sequences, positions of k-mers are more significant as compared to their semantics. Experimental results on a public benchmark dataset has proved that proposed feature representation approach significantly improves the performance of convolutional and recurrent neural network based approaches for the task of miRNA sub-cellular location prediction. In addition, experimental results prove that sequence to sequence models do not perform well for multi-label classification as they highly depend on label order information. Two simple models have significantly outperformed the performance of state-of-the-art MIRLocator approach based on sequence to sequence model. Considering the effectiveness of presented kmerRP2vec feature representation approach, we believe it can also be used to improve the performance of other DNA and RNA classification tasks. In future, we will assess the performance impact of proposed feature representation approach in multifarious DNA and RNA classification tasks and will also design robust classification model for miRNA subcelular localization.

## Footnotes

1 http://www.mirbase.org

